# Reporting quality of quantitative polymerase chain reaction (qPCR) methods in scientific publications

**DOI:** 10.1101/2024.12.04.626769

**Authors:** Natascha Drude, Camila Baselly, Małgorzata Anna Gazda, Jan-Niklas May, Lena Tienken, Parya Abbasi, Tracey Weissgerber, Steven Burgess

**Affiliations:** QUEST Center for Responsible Research, Berlin Institute of Health at Charité - Universitätsmedizin Berlin, Germany; Department of Biological Sciences, University of Montréal,1375 Avenue Thérèse-Lavoie-Roux, H3C 3J7 Montréal, Québec, Canada; Institute for Experimental Molecular Imaging, University Hospital RWTH Aachen, Germany; Medizinisches Kompetenzzentrum |c/o HCx Consulting GmbH | Brandenburg, Germany; 1. Department of Plant Science, University of Illinois Urbana Champaign. 2. Carl R Woese Institute for Genomic Biology, University of Illinois Urbana Champaign, United States of America

**Keywords:** qPCR, Reporting Quality of qPCR, Plant Science, Genetics, Reporting Standards, qRT-PCR, RT-qPCR, Quantitative reverse transcription polymerase chain reaction, (m)RNA quantification, meta-research, cross-sectional study

## Abstract

Reproducibility is a significant concern in scientific research and complex methods like quantitative polymerase chain reaction (qPCR) demand stringent reporting standards to ensure that the methods are reproducible, data are sound, and conclusions are trustworthy. Although the MIQE (Minimum Information for Publication of Quantitative Real-Time PCR Experiments) guidelines were introduced in 2009 to improve qPCR reporting, a 2013 study identified ongoing deficiencies that hinder reproducibility. To further investigate the transparency and completeness of qPCR reporting, we systematically assessed articles published in the top 20 journals in genetics and heredity (n=186) and plant sciences (n=246) that used qPCR. Our analysis revealed frequent omissions and inadequate specification of critical information necessary for evaluating and replicating qPCR experiments. RNA integrity, along with assessment methods and instruments used to assess it, are seldom reported. Although primer sequences are often disclosed, names and accession numbers of housekeeping genes are frequently omitted. Additionally, essential details about RNA extraction, RNA-to-cDNA conversion, and qPCR, such as kit names, catalog numbers, and reagent information, are often missing. Our findings underscore the urgent need for improved reporting practices in qPCR experiments, emphasizing quality controls, detailed descriptions of reagents and materials, and greater analytical transparency. Addressing these reporting deficiencies is crucial for enhancing the reproducibility and evaluating the trustworthiness of qPCR research. Potential solutions include encouraging authors to cite protocols published in online repositories, providing reporting templates, or developing automated tools to check reporting compliance.

## Introduction

Trust and the ability to build upon knowledge across different systems, designs, and disciplines are crucial for scientific advancement. The reproducibility crisis has become a prominent issue in scientific research, with the results of numerous studies across various fields failing to replicate^1–4^. This crisis underscores the importance of detailed and transparent methodological reporting, as inadequate reporting can obscure key details of the experimental design and execution, leading to irreproducible findings. Quantitative real-time PCR (qPCR) is widely used in molecular biology to measure RNA abundance in biological samples. qPCR measurements are sensitive to many factors (see Table 1), including the specificity of primers, the choice and stability of internal controls^5^ and reaction conditions, and the type of corrections performed by data processing software. Given the sensitivity and complexity of qPCR, detailed methods are essential to evaluate the trustworthiness of results and allow researchers to reproduce experiments. Herein we refer to methods reproducibility according to Godman et al. which “refers to the provision of enough detail about study procedures and data so the same procedures could, in theory or in actuality, be exactly repeated.”^3^

**Table 1:** Factors influencing the quality of the quantitative real-time PCR (qPCR) results.

| Component/<br>Stage | Potential Issues | Crucial items to report |
| --- | --- | --- |
| Reagents/<br>Materials | • Unspecific amplification of off-targets leads to inaccurate values.<br>• Spliceform specific amplification, leading to loss of information and inaccurate estimation.<br>• Primer dimers lead to false positive results.<br>• Unstable reference genes leading to incorrect values under experimental conditions<sup>6</sup> | • Primer binding site<br>• Primer concentration<br>• Reagents and enzymes<br>• Type of RT primer<sup>7</sup><br>• PCR machine<sup>8</sup><br>• Multiple reference genes<br>• Calculate reference gene stability<sup>6</sup> |
| Experimental<br>design | • Inconsistent reaction performance<sup>7,8</sup> leading to problems of reproducibility | • Annealing temperature<br>• Input RNA concentration |
| Quality assurance | • RNA degradation<sup>7</sup> leading to inaccurate quantification<br>• Presence of PCR inhibitors leads to inefficient amplification and distortion of values.<br>• Presence of gDNA contamination leads to overestimation of transcript abundance | • RNA integrity measurements<br>• RNA extraction method and kit<br>• Use of no-RT controls, or primer design |
| Data/ Analysis | • Different approaches to data analysis lead to different conclusions from the same data | • Setting a threshold for detection<br>• Software corrections and data processing procedures<br>• Calculation method<br>• Software<br>• Statistical methods<br>• Replicates, i.e. experimental unit |

Abbreviations: gDNA, (genomic DNA); RT, (Reverse Transcription)

To address concerns about poor reporting, the “Minimum Information for Publication of Quantitative Real-Time PCR Experiments” (MIQE) guidelines were introduced in 2009^9^. These guidelines outline the essential methodological details required to implement qPCR methods, critically evaluate them, and avoid drawing potentially misleading conclusions from qPCR data. Despite introducing these guidelines, surveys of papers published between 2009 and 2013 revealed widespread reporting deficiencies^10^. These studies showed that many articles failed to report critical information such as RNA integrity, primer sequences, and the specifics of the qPCR reagents and conditions used. Even though the MIQE guidelines are widely cited, with over 15,000 citations as of February 26, 2024, according to Google Scholar, recent reviews briefly assessed the current state of qPCR reporting practices and highlighted ongoing challenges^11–14^. An analysis of a small sample of 50 papers with qPCR data from 2023 showed that the guidelines are rarely referenced, and even when they are, often vital information is missing^15^. The complexity of the MIQE guidelines may discourage comprehensive adherence. As these guidelines are now more than 15 years old, there is a pressing need for updated assessments that employ a systematic, standardized approach to evaluate reporting quality.

In our study, we systematically examined qPCR reporting practices among articles published in the top 20 journals in genetics and heredity, and plant sciences. We evaluated the reporting of critical items such as primer sequences, housekeeping genes, normalization methods, and statistics (Figure 1). Our goal was to identify strengths and opportunities to improve reporting practices, thereby enhancing the reproducibility and integrity of qPCR research.

**Figure 1:**
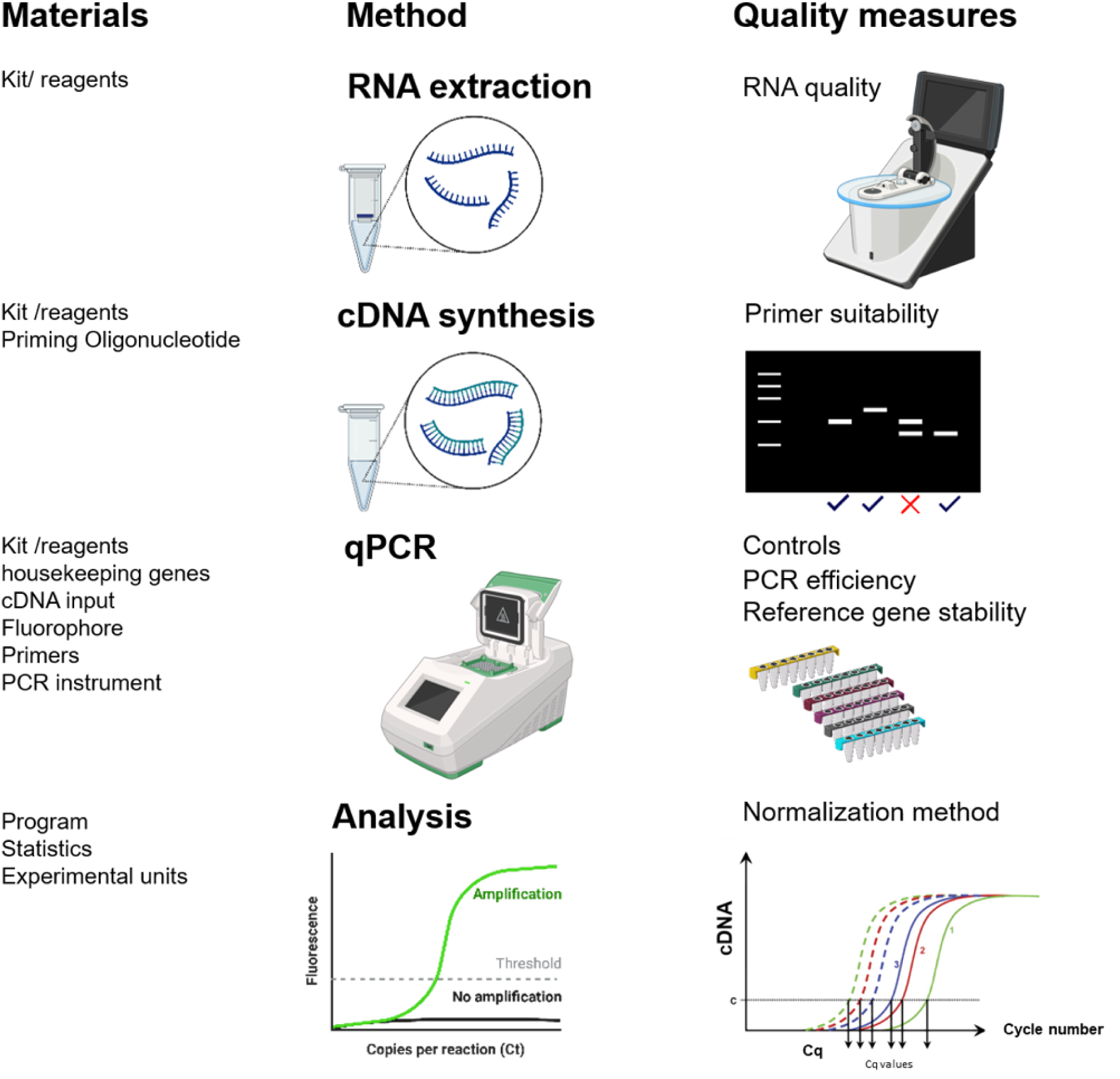
Overview of essential steps and materials in the qPCR workflow, including RNA extraction, cDNA synthesis, qPCR, and data analysis. Quality control measures, such as RNA integrity checks, normalization methods, and the use of appropriate kits, are highlighted to ensure reliable and reproducible results. Created with BioRender.com

## Methods

The abstraction protocol, data, and code for visualizations are available via the Open Science Framework^16^.

### Pre-registration

The study was pre-registered on the Open Science Framework (https://osf.io/9zp5m).

### Journal screening

The top 20 journals that published original research in each field of interest, namely genetics and heredity (Table S1), and plant science (Table S2), were identified by searching Journal Citation Reports. Journal lists, ordered by 2019 impact factor, were obtained for each field. Starting with the journal with the highest impact factor, the website for each journal was systematically examined. Journals that did not publish original research were excluded.

### Search strategy

Preliminary analyses revealed that the plant sciences journals published more papers per month than the genetics and heredity journals. Based on a random sample of 10 papers per journal, the plant sciences papers were also more likely to use qPCR. To compensate for this difference, we chose to examine papers published during a one-month period for plant sciences and a two-month period for genetics and heredity. Our goal was to obtain approximately 200 eligible articles per field.

Articles published in the top 20 genetics and heredity journals between September 1 and October 31, 2021 were identified using the following PubMed search: (“Nature genetics”[Journal] AND 53[Volume] AND (9[Issue] OR 10[Issue])) OR (“Genome research”[Journal] AND 31[Volume] AND (9[Issue] OR 10[Issue])) OR (“Molecular biology and evolution”[Journal] AND 38[Volume] AND ( 9[Issue] OR 10[Issue])) OR (“genome biology”[Journal] AND (“2021/09/01”[Date - Publication]: “2021/10/31”[Date - Publication])) OR (“genome medicine”[Journal] AND (“2021/09/01”[Date - Publication]: “2021/10/31”[Date - Publication])) OR (“American journal of human genetics”[Journal] AND 108[Volume] AND (9[Issue]OR 10[Issue])) OR (“Genes development”[Journal] AND 35[Volume] AND (17[Issue] OR 18[Issue] OR 19[Issue] OR 20[Issue])) OR (“Molecular therapy: the journal of the American Society of Gene Therapy”[Journal] AND 29[Volume] AND (9[Issue] OR 10[Issue])) OR (“Genetics in medicine: official journal of the American College of Medical Genetics”[Journal] AND 23[Volume] AND (9[Issue] OR 10[Issue])) OR (“Oncogene”[Journal] AND 40[Volume] AND (35[Issue] OR 36[Issue] OR 37[Issue] OR 38[Issue] OR 39[Issue] OR 40[Issue] OR 41[Issue] OR 42[Issue] OR 43[Issue])) OR (“Am J Med Genet C Semin Med Genet”[Journal] AND 187[Volume] AND 3[Issue]) OR ((“Genomics, proteomics bioinformatics”[Journal]) AND (19[Volume])) AND (5[Issue]) OR (“Genomics “[Journal] AND 113[Volume] AND 5[Issue]) OR (“molecular autism”[Journal] AND (“2021/09/01”[Date - Publication]: “2021/10/31”[Date - Publication])) OR (“Human genetics”[Journal] AND 140[Volume] AND (9[Issue] OR 10[Issue])) OR (“npj genomic medicine”[Journal] AND (“2021/09/01”[Date - Publication]: “2021/10/31”[Date - Publication])) OR (“Horticulture research”[Journal] AND (“2021/09/01”[Date - Publication]: “2021/10/31”[Date - Publication])) OR (“The CRISPR journal “[Journal] AND 4[Volume] AND 5[Issue]) OR (“PLoS genetics “[Journal] AND (“2021/09/01”[Date - Publication]: “2021/10/31”[Date - Publication])) OR (“Human molecular genetics”[Journal] AND 30 [Volume] AND (17[Issue] OR 18[Issue] OR 19[Issue])) Seventeen of the top 20 plant sciences journals were indexed in PubMed. Horticulture Research was on both the genetics and heredity and plant sciences lists, and was included in the genetics and heredity search above. Articles published in the remaining 16 journals in September 2021 were identified using the following PubMed search strategy: (“Nature plants”[Journal] AND 7[Volume] AND 9[Issue]) OR (“Molecular plant”[Journal] AND 14[Volume] AND 9[Issue]) OR (“The plant cell”[Journal] AND 33[Volume] AND 9[Issue]) OR (“The new phytologist”[Journal] AND 231[Volume] AND (5[Issue] OR 6[Issue])) OR (“Plant biotechnology journal “[Journal] AND 19[Volume] AND 9[Issue]) OR (“Plant physiology”[Journal] AND 187[Volume] AND 1[Issue]) OR (“Plant, cell environment”[Journal] AND 44[Volume] AND 9[Issue]) OR (“The Plant journal: for cell and molecular biology”[Journal] AND 107[Volume] AND (5[Issue] OR 6[Issue])) OR (“Journal of experimental botany”[Journal] AND 72[Volume] AND (17[Issue] OR 18[Issue])) OR (“Journal of integrative plant biology”[Journal] AND 63[Volume] AND 9[Issue]) OR (“TAG. Theoretical and applied genetics. Theoretische und angewandte Genetik”[Journal] AND 134[Volume] AND 9[Issue]) OR (“Frontiers in plant science”[Journal] AND (“2021/09/01”[Date - Publication]: “2021/9/30”[Date - Publication])) OR (“Molecular plant pathology”[Journal] AND 22[Volume] AND 9[Issue]) OR (“Phytomedicine: international journal of phytotherapy and phytopharmacology”[Journal] AND 90[Volume]) OR (“Physiologia plantarum”[Journal] AND 173[Volume] AND 2[Issue]) OR (“Plant cell physiology”[Journal] AND 62[Volume] AND 9[Issue])

The journal Preslia was excluded because it did not publish an issue in September 2021. Articles for the remaining two plant sciences journals, Journal of Ecology, and Environmental and Experimental Botany, were not available through PubMed; therefore the September 2021 issues of these journals were downloaded directly from the journal websites. All remaining journals were indexed in PubMed; therefore secondary searches using other search engines were not performed.

In accordance with the time periods for each field, articles published in Horticulture Research in September 2021 were included in datasets for both fields, whereas articles published in October 2021 were only included in the genetics and heredity dataset.

### Article Screening

Articles were screened to identify all full-length original research articles that used qPCR, focusing on applications looking at quantitative differences in gene expression. Studies that involved semi-quantitative qPCR analysis, droplet digital PCR, Northern Blot analysis, copy variant number analysis on genomic DNA were therefore excluded. Articles that were not full-length original research were excluded. Screening was performed in Rayyan (RRID:SCR017584) by two independent reviewers (SJB, MAG). Disagreements were resolved by consensus.

### Data Abstraction

Data were abstracted by a group of six independent reviewers in rotating pairs (LT, JNM, MAG, PA, NID, SB). Abstraction was performed online using Microsoft Forms (for abstraction form see OSF^16^), following the pre-registered protocol. Disagreements were resolved by consensus, first by the abstracting duo and if needed by the whole consortium. Each article was assessed to determine whether the following information was reported:

1. **Location of qPCR methods:** Where did authors describe qPCR methods and/or materials (e.g., methods section, supplemental files, methodological shortcut citation^17^)? Authors use a methodological shortcut citation when they cite another resource, instead of fully describing the method.
2. **Guideline use:** Did the paper cite the Minimum Information for Publication of Quantitative Real-time PCR Experiments (MIQE^9^)?
3. **RNA details:** Was the RNA extraction method specified? Did the authors report the RNA integrity/quality and specify the instrument used for this measurement?
4. **Reverse transcriptase protocol:** Did the authors report the kit name, the priming oligonucleotide, and additional details or modifications of kit procedures?
5. **PCR protocol**: Did the authors report complete reaction conditions, the reaction volume, the cDNA input, the buffer or kit name, the PCR instrument, and PCR efficiency?
6. **Primer information and reference genes:** Did the authors report the primer sequences, their target sequence accession number, the specificity of primers, and the housekeeping genes? Did the authors report reference gene stability (M value)?
7. **Normalization method:** Did the authors report the normalization method (double delta Cq (ddCq) (Cq = quantification cycle), or normalized relative quantity (NRQ))?
8. **Fluorophore:** Did the authors report whether they used an intercalated dye (e.g., SYBR/TB Green) or an oligonucleotide fluorescent reporter (e.g., Taqman)?
9. **Cq values (Control):** Did the authors report the Cq values of an NTC/NAC (no template control or no amplification control) and the Cq values of an NRT/MRT (negative reverse transcriptase or minus reverse transcriptase) control?
10. **qPCR software:** Did the authors report what qPCR analysis program (source, version) was used?
11. **Statistical method:** Did the authors report the methods used to assess statistical significance?
12. **Replication number:** Did the authors specify the number of experimental units for each group?

### Changes to the pre-registered abstraction protocol

Two variables were removed from the protocol because they could not be assessed reliably. These items examined whether the authors defined control and experimental groups, and the experimental unit. Answer options for the question on fluorophores were renamed to improve clarity. Answer options for questions on RNA extraction kits, the instrument used to assess RNA integrity, and the name of the real-time PCR kit were expanded to capture more information. Detailed changes are available in the “Protocol_changes” document on <u>OSF</u>.

### Data Visualization and Analysis

We calculated the percentage of papers in each field that reported each item. Figures were created using the R Project for Statistical Computing (RRID:SCR_001905) Version 4.2.2. Data and code are available on the Open Science Framework^16^. This was an exploratory, descriptive study examining qPCR reporting practices in two fields; therefore, formal statistical analyses were not performed.

## Results

The study included the top 20 genetics and heredity, and plant sciences, journals based on their 2019 impact factor. Six genetics and heredity journals and six plant science journals were excluded because they did not publish original research (Figure 2). 181 articles from genetics and heredity and 189 articles from plant science were excluded, as they were not original research articles. Another 375 articles from plant science and 331 from genetics and heredity were excluded based on our predefined inclusion and exclusion criteria.

**Figure 2:**
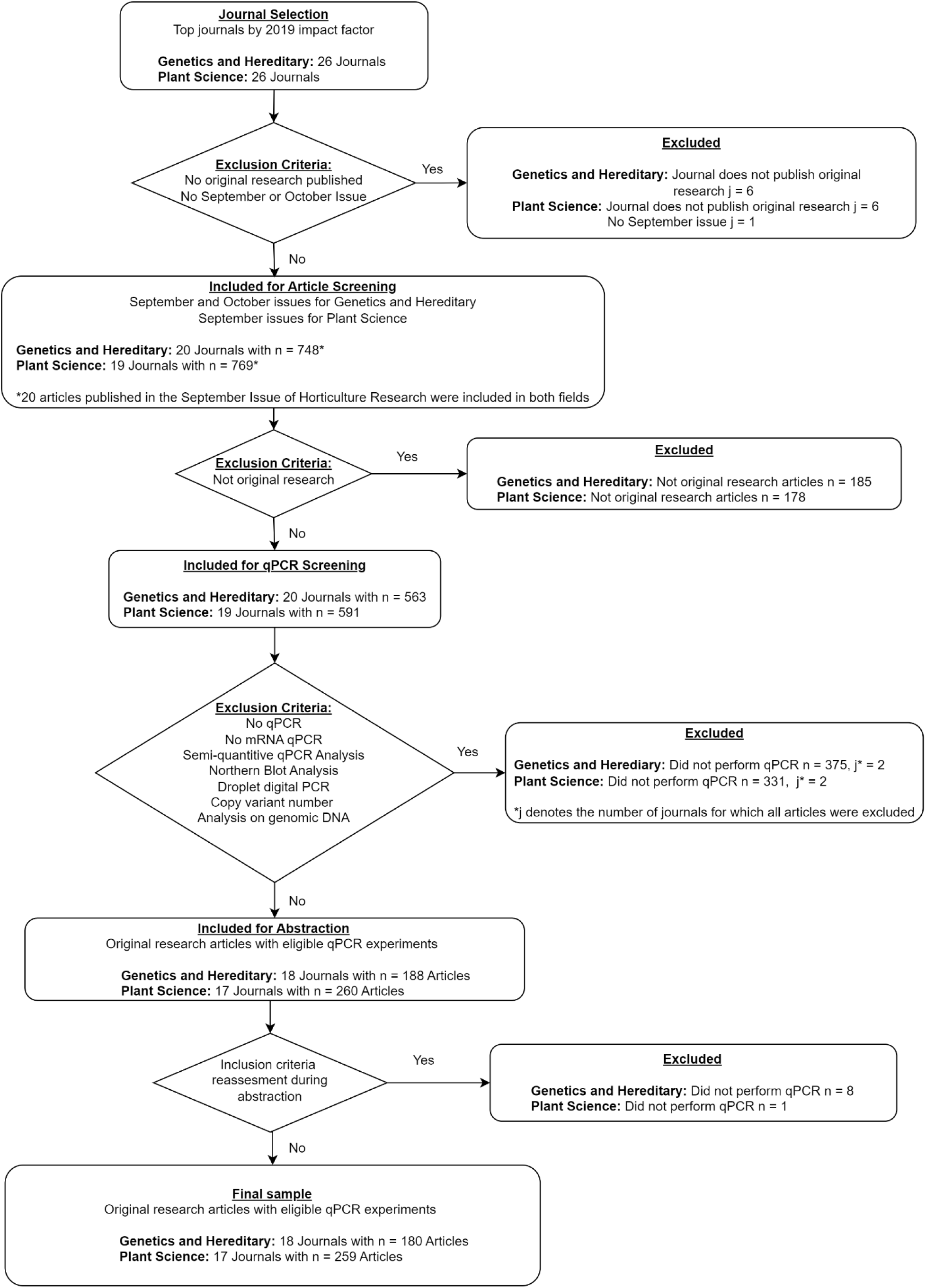
Study flowchart. This flow chart illustrates the journal and article screening process and shows the number of observations excluded and reasons for exclusion at each phase of screening. Abbreviations: j = number of journals; n = number of articles. Data is available at the following OSF repository^16^: https://osf.io/2ps43/ in the ‘Data’ folder.

### Reporting of qPCR Information

qPCR methods were described in various locations: in the main manuscript (98.8%), supplementary material (86.7%), repository (0%), or via shortcut citations of other sources (30.1%). When assessing reporting for the remaining items, we only considered methods reported in the main article (including figures) and supplementary materials. We did not follow any shortcut citations, nor did we extract information from linked resources like repositories.

## 1. Reporting of materials, reagents, and kits

The majority of articles provided information on the RNA extraction kit or reagent, the RT or PCR kit, the name of the housekeeping gene(s), the primer sequences, and the cycler used (Figure 3). Despite the generally good reporting of primary materials, there were significant gaps in the reporting of specific details. Most articles failed to provide exact information such as target sequence accession numbers (only 31% in plant science and 14% in genetics) or catalog numbers (8% in plant science and 14% in genetics). Additionally, while 88% of genetics and heredity articles and 92% of plant science articles reported the primer sequences, only 9-10% of articles in both fields reported the priming oligonucleotide.

**Figure 3:**
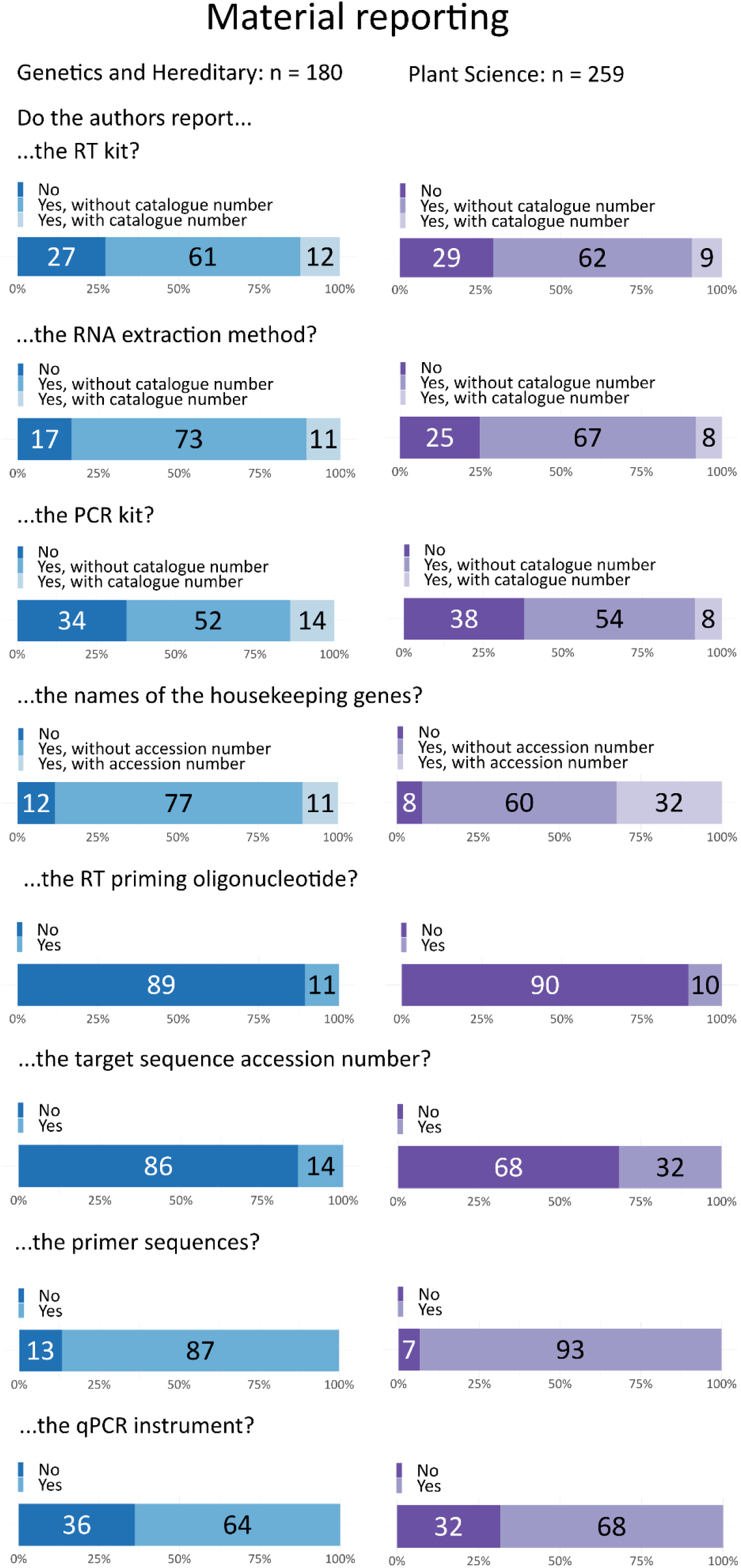
Reporting of materials and instruments for qPCR method reproducibility. Both fields consistently report essential components such as reagents or kits used for RNA extraction, reverse transcription, PCR, as well as details of the cycler, housekeeping gene, and primer sequences. However, most papers failed to report accession numbers and priming oligonucleotides.

## 2. Method reporting

We evaluated two key aspects of qPCR methods reporting: the information needed to reproduce an experiment and the quality aspects that ensure the experiment was conducted with high precision. Methodological details needed to reproduce an experiment included annealing temperatures, cycling protocols, the concentration or mass of input materials, and other specific settings. Quality assurance measures included primer specificity, RNA integrity, and other critical factors that contribute to the reliability and accuracy of the qPCR results.

### 2.1 Conduct of experiments (methods reproducibility)

Our study revealed that most articles fail to provide comprehensive information on methodological details necessary for reproducing a method or result - although for procedures such as RNA extraction, cDNA synthesis, or PCR methods, if the authors stated that the procedure “was performed according to the manufacturer’s instructions,” we granted the authors the benefit of the doubt and counted this as adequate reporting. “Unclear” cases are those where the kit is mentioned and parts of the methods are described, but it is not stated if any modifications were made. However, even with this leniency, over 98% of papers in both fields failed to provide sufficient information on any modifications or the actual procedures used (Figure 4). Most manuscripts did not provide details on the annealing temperature (genetics: 80%, plant sciences: 72%) or the overall cycling protocols.

**Figure 4:**
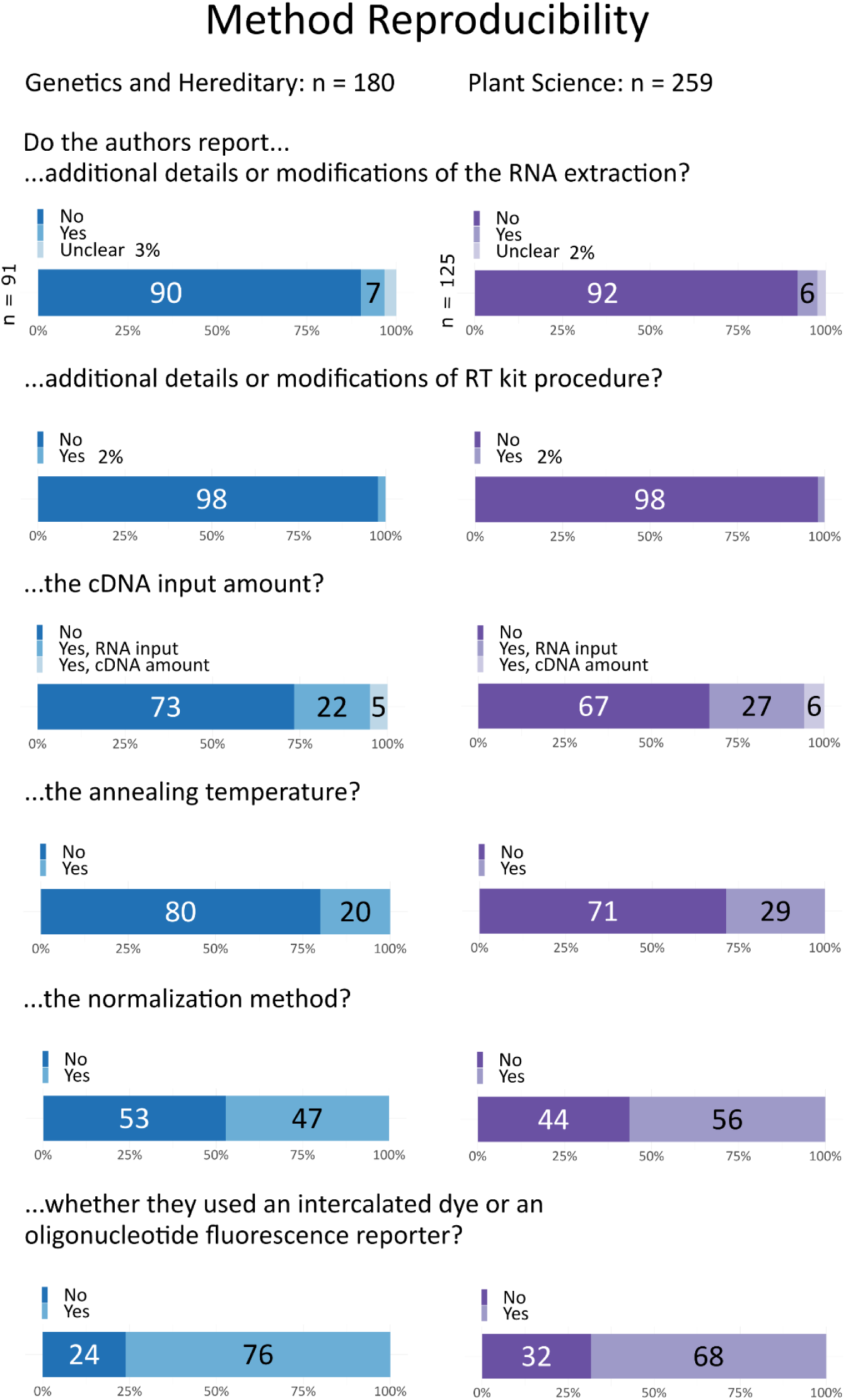
Reporting of methods related to procedures, concentrations, and measurement settings. Both fields lack sufficient details required for fully reproducing the measurements.

The input amount of cDNA was only reported in 27% of genetics papers (with 22% specifying the RNA input and 5% the actual cDNA amount) and in 34% of plant science publications (with 28% specifying the RNA input and 6% the cDNA input). Importantly, we only considered the input amount as reported if it was given as a concentration or mass; descriptions such as “5 µL were used” without specifying the concentration were considered as “not reported.” One issue with only reporting the RNA input is that the enzyme reverse transcriptase can introduce a bias, resulting in cDNA not being proportional to the RNA input. This can be investigated by generating standard curves of target and control assays.

The type of fluorophore used, such as TaqMan or SYBR Green, was provided in most articles (genetics: 76%, plant science: 67%). If SYBR Green was used, we looked into critical controls like no template controls.

The name of the normalization method was reported in 47% of genetics articles and 57% of plant science articles. However, details were often relegated to a shortcut citation that we did not abstract or check.

### 2.2 Quality assurance measures

Information on RNA integrity, such as RNA Integrity Number (RIN) or RNA Quality Indicator (RQI) values, or integrity assessment via RNA gel, was rarely provided (genetics: 7%, plant sciences: 10%, Figure 5). Often, this was accompanied only by a qualitative statement that the assessment was done, without any values or supplementary materials like images of the RNA gel. When authors did provide information on RNA integrity, it was often unclear what device was used for the assessment. Manuscripts that included RNA sequencing frequently provided details on RNA integrity, but it was often unclear if the same samples and procedures were used for the qPCR samples. In cases where the instrument was mentioned, details typically included the device manufacturer.

**Figure 5:**
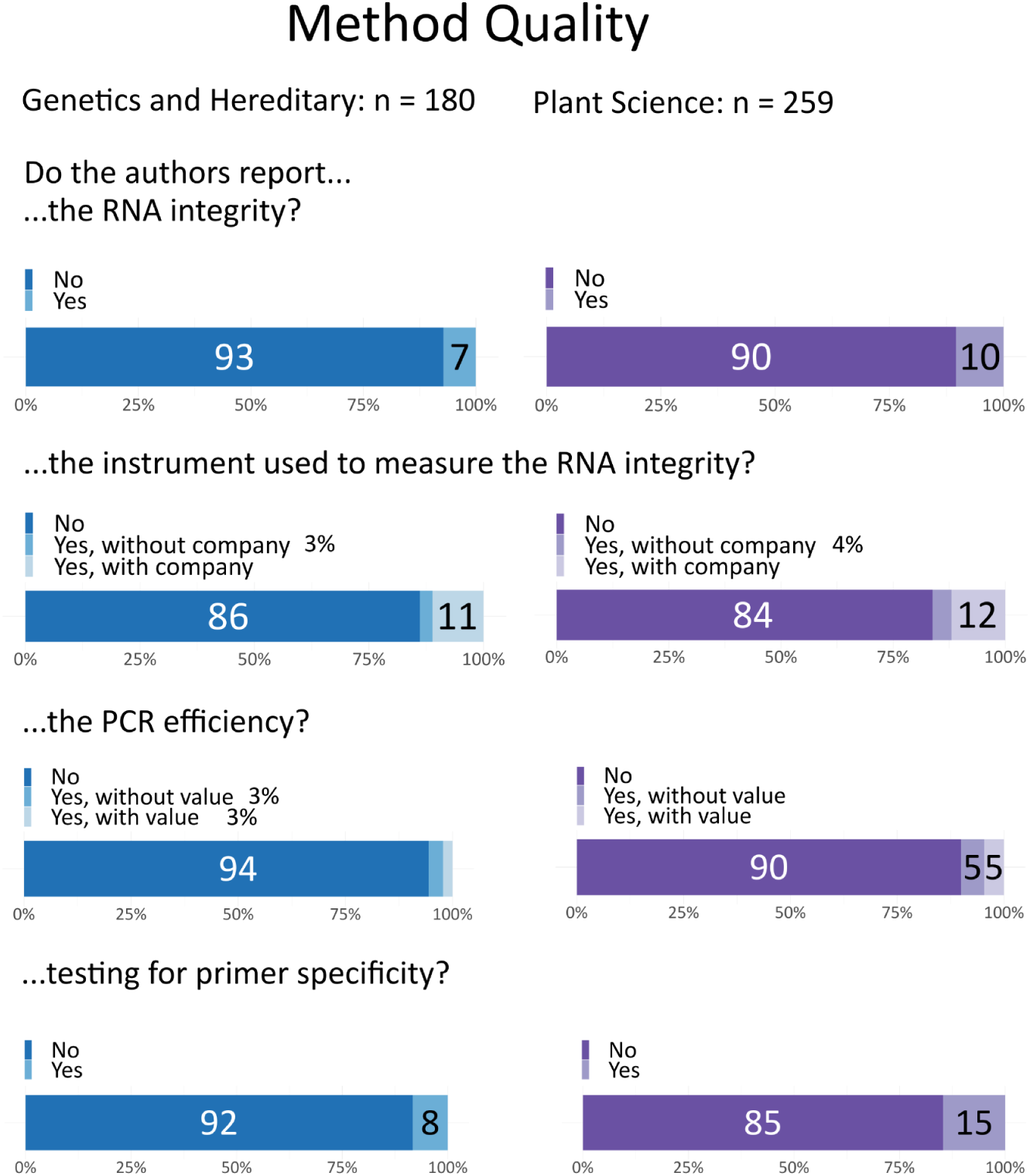
Reporting of quality measures during qPCR.

We also examined the specificity of primers, which can be assessed using a DNA agarose gel or melt curve analysis. We classified this item as reported if the authors stated that these checks were done. Primer specificity was rarely reported (genetics: 8%, plant sciences: 13%).

For fluorophores like SYBR Green, we checked if the article provided the Cq values of a no template control (NTC) or no amplification control (NAC) (Figure S1). Additionally, we looked for the Cq values of a negative reverse transcriptase (NRT) or minus reverse transcriptase (MRT) control. When available, this information was often hidden in graphs or tables, as well as in the main text or supplementary materials, with its location varying from paper to paper.

Regarding PCR efficiency, which is typically calculated from the slope of a standard curve and provided as the correlation coefficient (R²), this was reported in only 5% of genetics articles and 8% of plant science articles, with only half of these providing actual values. Gene stability, assessed via average expression stability (M-value), was not reported (0.4%) in either field (Figure S2).

Overall, our analysis revealed substantial reporting deficiencies in qPCR quality measures. Poor reporting does not necessarily mean that these assessments were not performed. However, the lack of detailed reporting makes it difficult for readers to judge the trustworthiness, robustness, and reliability of qPCR results.

## 3. Statistical reporting

Reviewers were often unable to determine which statistical tests were used for qPCR data, as statistical methods were typically described in a separate paragraph in the methods section. These general descriptions mentioned multiple tests, without specifying which test was used for which experiment.

Initially, we sought to determine whether authors clearly distinguished between biological replicates and technical replicates. We found that authors often used terms like “triplicates” or “independent replicates” without clarifying whether these were biological or technical replicates. The definitions of technical and biological replicates varied widely and were often not clearly delineated by the authors. Biological replicates usually refer to independently derived samples that represent biological variation (e.g., samples from different plants), however, samples may also be considered as biological replicates if they are from the same organism, but variation has been artificially introduced to each batch. Technical replicates can involve repeated measurements of the same sample to assess the precision of the technique, but can also be defined as different batches from the same sample. Given the poor reporting, combined with the potential for bias and subjectivity in interpreting these definitions, we decided that it was not possible to extract this information in a consistent, high-quality manner. Instead, we focused on whether the experimental unit or sample size was reported. The experimental unit is the entity that is randomly and independently assigned to experimental conditions. This is equal to the sample size (n)^18^. Seventy-five percent of the genetics articles and 84% of the plant science articles provided this information. This includes any type of replicate, however, and essential details about whether replicates were independent was often missing.

## Discussion

Our findings indicate significant gaps in the reporting of methodological details, which are crucial for the reproducibility and accurate interpretation of qPCR experiments. More detailed reporting is needed to improve the transparency and reliability in qPCR research. While most papers reported primer sequences (>88%) and provided the names of reference genes, evidence for validation of assay efficacy was rare. This information is not needed for reproducibility, but is critical for assessing the validity of the results. Most papers omit transcript accession numbers, placing the responsibility of verifying isoform and spliceform specificity on the readers. Validated primer pairs are essential for generating robust qPCR results, as various factors can influence assay efficacy^19^. Key considerations include:

1. Primer specificity for the target transcript: Primer specificity is critical to avoid noise from cross-amplification. Given that many transcripts undergo alternative splicing, it is important to report the precise location of primer binding sites along with corresponding transcript identifiers. In organisms that have undergone whole genome duplication, or contain gene duplicates, designing primers for specific homologs can be challenging^20^.
2. Formation of ‘primer-dimers’: These short unspecific amplification products are formed by complementary pairing of a pair of primers at the 3’ end allowing DNA polymerase to extend. In some instances, a pair of primers bind non-specifically in close proximity to an alternate target, generating short DNA fragments^21^. Both these processes will lead to overestimation of transcript abundance.
3. Hairpins: Unlike textbook depictions, single-stranded DNA frequently exists in stable secondary structures, called hairpins. If a primer forms a hairpin or is designed against a folded region, this structure must be broken before the primer can bind.^21^
4. Primer annealing temperature (Ta): This is the temperature at which a maximum amount of primer is bound to a template. A variety of algorithms have been developed to assist with primer design, such as PrimerSeq^22^, Visual-OMP^21^, and PrimerBLAST^23^. The secondary structure of amplicons can be tested for hairpins with mfold^24^ or UNAfold^25^.

While *in silico* analysis can act as an initial screen to increase the chance of a successful assay, the efficiency of primers needs to be empirically assessed^26^. This is important because some algorithms do not calculate important variables, such as Ta, or use incorrect parameters for a particular reaction setup^19^. Over 90% of papers did not mention attempts to test for efficiency or specificity (Figure 5), and a further 3-5% mentioned specificity was tested without providing evidence. In addition, assays can behave differently depending on the reagents^27^, quantity of input RNA^7^, primer concentration, annealing temperature^19^ or qPCR machine^8^. Further, the choice of reverse transcriptase can result in up to 100-fold variation in cDNA yields^28^, and the priming strategy (random hexamer, gene-specific primers, oligo dT) can also influence results, with up to 24-fold difference reported depending on the gene analyzed^7^. Reporting of assay conditions was also poor, with the majority of papers failing to provide details about input nucleic acid concentrations (>66%) or the annealing temperature used in qPCR programs (>69%). While the reporting of RT kits (>70%) and qPCR machines (>64%) used was generally better, these were often unaccompanied by accession numbers which has the potential to result in incorrect choice of product during efforts to reproduce experiments. An analysis of qPCR reporting in plant science journals in 2008 found that approximately 3% of papers used validated reference genes^29^, which is in general agreement with our findings, suggesting repeated calls for improvements in reporting^30–34^ have had little impact. In summary, due to inadequate reporting, it is impossible to determine whether >90% of qPCR results are reliable and less than half of experiments would be able to be repeated under the same conditions used in manuscripts. Rigorously conducted research can be reported poorly; therefore it is not possible to determine what proportion of conclusions might be affected.

qPCR results are normalized to reduce sample-to-sample variation, and while the quantity of input RNA and sample size are sometimes used, the amplification of an internal standard is the most common^35^. So-called reference or ‘housekeeping’ genes are chosen due to their ubiquitous expression, relative stability, and high abundance^5^. Unfortunately, most reference genes are used without validation^6,35^, despite extensive evidence that expression is constant under some conditions and in some tissue types, but can be highly variable under others^36–40^. This issue can be compounded by the use of a single reference gene, an approach which is estimated to result in erroneous normalization of >3 fold in 25% of cases^6^. As a result, inappropriate choice of internal standards can alter findings^41^. This has led to recommendations to use 3 or more reference genes^6^ and the development of statistical algorithms to assess the stability of their expression over experimental conditions, such as geNorm^42^, NormFinder^43^, BestKeeper^44^, and RefFinder^45^. We did not record the number of reference genes used in experiments here, but we found no evidence in our sample of any efforts to test, or report, the stability of reference genes, raising serious questions about the reliability of presented data.

Even if primers are correctly validated, experimental noise can be introduced by sample processing, and different results can be obtained depending on the RNA storage and extraction method^46,47^. Co-purification of inhibitors during processing reduces the efficiency of reverse transcription and PCR^48^, and RNA quality can influence results^49–51^, as there is a high correlation (R^2^>0.8) between RNA integrity and measured transcript abundance^52^ which cannot be corrected by normalizing to reference genes due to differences in transcript degradation rates^53^. Another consideration is that genomic DNA contamination is common in purified RNA, even after treatment with DNAse I. Bustin found that RNA constituted, on average, 50-80% of nucleic acid isolated using a commercial kit, with values ranging from almost pure RNA, to almost pure DNA, depending on the tissue of origin and the experimentalist^54^. This can be a problem for accurate quantification, particularly in the case of low abundance transcripts.

In addition, it is common for qPCR and RNA sequencing experiments to be described within the same paper. However, quality control measures for RNA are often reported only in the context of RNA-seq, while similar details are omitted for qPCR. This creates uncertainty regarding whether the same RNA, with the same quality metrics, was used for both RNA-seq and reverse transcription in qPCR. This issue is further complicated when slightly different RNA extraction methods are employed for each technique, making it difficult to assess consistency and data reliability.

qPCR assays can be made insensitive to gDNA contamination by designing primers which span exon junctions, but this is not always possible. Genomic DNA contamination is therefore typically assessed through analysis of “no-RT (NRT)” reactions, where samples are processed as normal, however, the enzyme is omitted from the reverse transcriptase step. After comparing experimental and NRT samples, an arbitrary threshold of <3% gDNA contamination is typically deemed acceptable^9^. As with checks for reference gene stability, we found no evidence for the use of NRT controls in our sample. While most papers reported some information about RNA extraction methods (80-85%), only a small percentage reported information about RNA integrity (11%). As checking whether primers were designed over exon boundaries was beyond the scope of this study, the extent to which RNA processing or gDNA contamination impacts the interpretation of qPCR studies is unclear.

Once an experimental run is complete, raw fluorescence data must be processed to determine target transcript abundance. The cycle quantification (Cq) value, which is sometimes referred to as cycle threshold (Ct) or crossing point (Cp), is the point at which measurable fluorescence rises above a background. The Cq value is typically determined by a ‘fit-point’, ‘cycle threshold’^55^ or ‘second derivative maximum (SDM)^56^ method. This value is directly related to the starting concentration of a target, and differences in Cq values between samples for a given primer pair are used to calculate the relative abundance of a transcript^5,57^. However, Cq values are sensitive to the calculation method and can be influenced by the PCR efficiency in addition to experimental artifacts (as reviewed by Ruiz-Villalba et al.^58^). Once Cq values are calculated, several different equations can be used to determine the relative normalized quantity (NRQ) of a target. One of the simplest is the ‘ddCT’ method, whereby the Cq values of reference and target genes are compared between samples, assuming a 100% PCR efficiency^57^. Pfaffl updated this approach taking into account the difference in amplification efficiencies between reference and target genes^59^, but this method was further modified^60^. All methods and equations are in use, and the chosen equation must be reported because this can influence experimental outcomes. In our sample, only 44-54% of papers report the exact method used to calculate gene expression.

A further complication is that qPCR machines come with their own analysis software which can apply different corrections. Several third party algorithms, including REST^61^ and qBase^60^ standard format for qPCR data, known as Real-Time PCR Data Markup Language (RDML)^6,62,63^. We found no mention of RDML file format in our sample and 48-59% of papers did not report which software was used to perform calculations (Figure 6).

**Figure 6:**
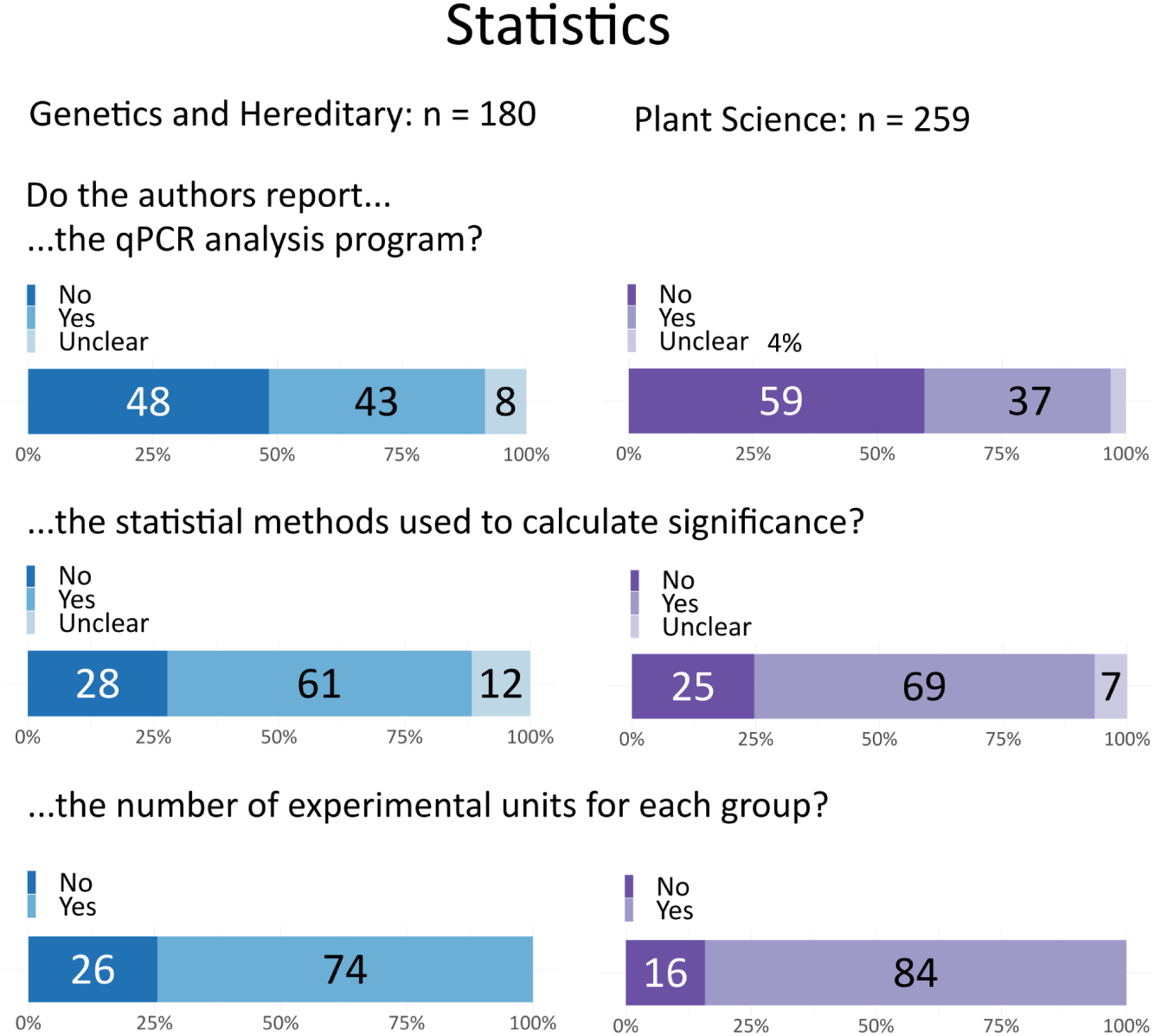
Reporting of analysis. The number of experimental units was found in most papers, whereas the statistical methods and qPCR analysis program were less commonly reported.

A previous analysis comparing papers between 2009-11 and 2012-13 suggested a slight, but significant, improvement in the quality of reporting following the release of the MIQE guidelines^64^. Our data based on publications from 2021 indicate that further progress is needed.

### Solutions

The persistence of poor methodological reporting for qPCR experiments in studies published 12 years after the release of the MIQE guidelines highlights the need to implement effective solutions. The precise causes of reporting issues are unclear but could range from overly stringent reporting standards to a lack of awareness around the limitations and pitfalls of a widely used technique. One solution may be to update and simplify the reporting guidelines.

The comprehensive MIQE guideline includes 85 elements. These are subdivided into 57 “essential” elements, which are crucial for reproducibility and should be reported by all authors, and 28 “desirable” elements, which authors are strongly encouraged to report. A similar approach of distinguishing between “essential” and “recommended” elements was recently used to redesign and simplify the ARRIVE (Animal Research: Reporting of In Vivo Experiments)^65^ guidelines for reporting preclinical animal studies after research suggested that the 2010 ARRIVE guidelines failed to substantially improve reporting quality^63^. Simplification has been attempted with the MIQE précis, or absolute minimum standard, which reduced the number of required elements to report to 29^66^. Further research is needed to determine whether this improved reporting. An updated MIQE guideline might also include reporting templates or training materials to illustrate good reporting for each item, building on existing efforts^67^.

A second solution may be to encourage authors to share reusable step-by-step qPCR protocols on open-access protocol repositories^68^ and cite these protocols in their papers. The general text descriptions found in the methods sections of papers are often missing essential details^69^. Step-by-step descriptions of procedures are often more useful to readers who want to implement a method^70^ and would make it easier to capture essential details required by the MIQE guidelines. Additionally, journal word count limitations may also play a significant role, as trimming methods sections for brevity might seem less daunting to authors than reducing the length of results or discussion sections. Providing external repositories for detailed protocols could alleviate this pressure by allowing authors to keep methods concise while ensuring all critical details remain accessible to readers. Authors who choose repositories that allow versioning can also update their protocol as it evolves while citing the version of the protocol used in a specific study. Sharing step-by-step protocols is a key recommendation for researchers in the European Commission PRO-MaP (Promoting Reusable and Open Methods and Protocols) report^70^. Using and maintaining updated protocols also benefits researchers by making it easier to standardize procedures within and across research groups, train new personnel, and locate the version of the protocol used in past studies.

A third consideration is whether improved training for researchers performing qPCR is needed. It is unclear whether the lack of reporting is due to overly burdensome standards, a lack of awareness about reporting requirements, or insufficient understanding of the potential pitfalls of performing qPCR.

Fourth, the stakeholders within the research community can work together to develop more efficient systems for checking reporting. Enforcement of standards is ultimately the responsibility of journal editors and reviewers, however, checking compliance with the guidelines is time-consuming. To reduce the burden on screening, one solution could be to develop automated tools that scan through manuscripts before submission and highlight areas where greater detail is required^71^. The commercial tool SciScore (https://www.sciscore.com/), for example, is designed to check items on the MDAR checklist^72^, and open source tools have been used to check other aspects of reporting^73^. The scattering of qPCR methods across different sections of the paper, including figure legends, methods sections, and supplementary data, would complicate these efforts. Supplemental files are especially challenging, due to the variety of file types and archiving systems used by different publishers.

### Limitations

Results of this study may not be generalizable to studies published in fields other than genetics and heredity and plant sciences, studies that are not published in English, or studies published in other journals. Some authors used shortcut citations, instead of fully describing their methods in the paper. We did not have the resources to assess shortcut citations to determine whether methodological details that were missing from the paper were provided in the cited resource. While shortcut citations can provide important methodological details, problems that one might encounter include difficulty identifying or accessing the cited resource, difficulty finding the cited method within the cited resource, an inadequate description of the cited method, or chains of shortcut citations^17^.

In many instances we were unable to assess whether researchers actually performed appropriate validation experiments, for example relating to primer specificity and amplification efficiency. In this study we accepted statements that said that validation was done, however, these statements often lacked supporting evidence. This may systematically bias our data towards better reporting.

Likewise, we recorded whether statistical methods and comparative groups were reported, but we did not assess whether the types of comparisons or methods used were appropriate. Plant sciences studies investigating the impact of transgene expression, for example, sometimes compare transgene expression between a transformed and a non-transformed organism. This comparison is inappropriate as the transgene is absent in the non-transformed organism. Similarly, papers sometimes compare relative expression between different genes. This practice should be avoided due to variations in primer efficiency, reverse transcription efficiency, amplicon length, and PCR efficiency.

While the MIQE guidelines are highly cited, only two papers in our sample (n=439) referenced the guidelines. We were unable to determine whether reporting is better among studies that cite the guidelines.

## Conclusion

In line with studies conducted 10-15 years ago, we found that the reporting of qPCR methods remains generally poor. Most manuscripts are missing essential information needed to implement the method, assess data quality, and evaluate the paper’s conclusions. Rigorously conducted research may be reported poorly; therefore, we were not able to determine how often methodological problems may have altered study conclusions. These results suggest that reporting guidelines alone are insufficient to ensure accurate reporting of qPCR methods. The research community needs to develop and test new solutions. This might include enhanced training, reporting templates, a simplified set of requirements, encouraging authors to share step-by-step qPCR protocols, or the use of automated tools to screen for compliance. Collectively, these efforts aim to support transparent, open, and reproducible science while also minimizing the waste of time, materials, and funding caused by irreproducible or incomplete reporting.

## CREDIT Authorship Statement

Conceptualization: Natascha Drude, Steven Burgess

Methodology: Natascha Drude, Steven Burgess, Małgorzata Anna Gazda, Tracey Weissgerber Investigation: Natascha Drude, Małgorzata Anna Gazda, Jan-Niklas May, Parya Abbasi, Lena Tienken, Steven Burgess

Data curation: Małgorzata Anna Gazda, Camila Baselly, Steven Burgess, Natascha Drude, Tracey Weissgerber

Formal analysis: Camila Baselly Project administration: Camila Baselly

Supervision: Natascha Drude

Validation: Camila Baselly, Tracey Weissgerber, Natascha Drude Visualization: Camila Baselly, Natascha Drude, Tracey Weissgerber

Writing – original draft: Natascha Drude, Steven Burgess, Tracey Weissgerber

Writing – reviewing and editing: Steven Burgess, Natascha Drude, Jan-Niklas May, Małgorzata Anna Gazda, Lena Tienken, Camila Baselly, Parya Abbasi, Tracey Weissgerber

## Acknowledgment

We would like to thank Maren Hülsemann for her invaluable assistance with the first training sets, and Constance Holman for her support during the initial screening of original research articles.

## References

1. Errington, T. M., Denis, A., Perfito, N., Iorns, E. & Nosek, B. A. Challenges for assessing replicability in preclinical cancer biology. eLife 10, e67995 (2021).

2. Errington, T. M. et al. Investigating the replicability of preclinical cancer biology. Elife 10, e71601 (2021).

3. Goodman, S. N., Fanelli, D. & Ioannidis, J. P. What does research reproducibility mean? Sci. Transl. Med. 8, 341ps12–341ps12 (2016).

4. Amaral, O. B., Neves, K., Wasilewska-Sampaio, A. P. & Carneiro, C. F. The Brazilian Reproducibility Initiative. eLife 8, e41602 (2019).

5. Thellin, O. et al. Housekeeping genes as internal standards: use and limits. J. Biotechnol. 75, 291–295 (1999).

6. Vandesompele, J. et al. Accurate normalization of real-time quantitative RT-PCR data by geometric averaging of multiple internal control genes. Genome Biol. 3, research0034.1 (2002).

7. Ståhlberg, A., Håkansson, J., Xian, X., Semb, H. & Kubista, M. Properties of the Reverse Transcription Reaction in mRNA Quantification. Clin. Chem. 50, 509–515 (2004).

8. Lu, S., Smith, A. P., Moore, D. & Lee, N. M. Different real-time PCR systems yield different gene expression values. Mol. Cell. Probes 24, 315–320 (2010).

9. Bustin, S. A., et al. The MIQE Guidelines: M inimum I nformation for Publication of Q uantitative Real-Time PCR E xperiments. (2009).

10. Taylor, S. C. & Mrkusich, E. M. The state of RT-quantitative PCR: firsthand observations of implementation of minimum information for the publication of quantitative real-time PCR experiments (MIQE). J. Mol. Microbiol. Biotechnol. 24, 46–52 (2014).

11. Abdel Nour, A. M., Nemer, G. & Khalil, A. The MIQE Guidelines’ tenth anniversary: The good and bad students. Gene Rep. 19, 100630 (2020).

12. Bustin, S. A. Improving the quality of quantitative polymerase chain reaction experiments: 15 years of MIQE. Mol. Aspects Med. 96, 101249 (2024).

13. Shipley, G. The MIQE guidelines uncloaked. PCR Troubl. Optim. Essent. Guide 151–165 (2011).

14. Borchardt, M. A. et al. The Environmental Microbiology Minimum Information (EMMI) Guidelines: qPCR and dPCR Quality and Reporting for Environmental Microbiology. Environ. Sci. Technol. 55, 10210–10223 (2021).

15. Bustin, S. A. Improving the quality of quantitative polymerase chain reaction experiments: 15 years of MIQE. Mol. Aspects Med. 96, 101249 (2024).

16. Heinrich, C. V.-Q. B. et al. Quality of reporting of quantitative polymerase chain reaction (qPCR) in scientific publications. (2024) doi:10.17605/OSF.IO/2PS43.

17. Standvoss, K. et al. Shortcut citations in the methods section: Frequency, problems, and strategies for responsible reuse. PLOS Biol. 22, e3002562 (2024).

18. Lazic, S. E., Clarke-Williams, C. J. & Munafò, M. R. What exactly is ‘N’ in cell culture and animal experiments? PLOS Biol. 16, e2005282 (2018).

19. Bustin, S. & Huggett, J. qPCR primer design revisited. Biomol. Detect. Quantif. 14, 19–28 (2017).

20. Zhao, F. et al. An optimized protocol for stepwise optimization of real-time RT-PCR analysis. Hortic. Res. 8, 179 (2021).

21. SantaLucia, J. Physical Principles and Visual-OMP Software for Optimal PCR Design. in PCR Primer Design (ed. Yuryev, A.) 3–33 (Humana Press, Totowa, NJ, 2007). doi:10.1007/978-1-59745-528-2_1.

22. Tokheim, C., Park, J. W. & Xing, Y. PrimerSeq: design and visualization of RT-PCR primers for alternative splicing using RNA-seq data. Genomics Proteomics Bioinformatics 12, 105–109 (2014).

23. Ye, J. et al. Primer-BLAST: A tool to design target-specific primers for polymerase chain reaction. BMC Bioinformatics 13, 134 (2012).

24. Zuker, M. Mfold web server for nucleic acid folding and hybridization prediction. Nucleic Acids Res. 31, 3406–3415 (2003).

25. Markham, N. R. & Zuker, M. UNAFold. in Bioinformatics: Structure, Function and Applications (ed. Keith, J. M.) 3–31 (Humana Press, Totowa, NJ, 2008). doi:10.1007/978-1-60327-429-6_1.

26. Svec, D., Tichopad, A., Novosadova, V., Pfaffl, M. W. & Kubista, M. How good is a PCR efficiency estimate: Recommendations for precise and robust qPCR efficiency assessments. Biomol. Detect. Quantif. 3, 9–16 (2015).

27. Buzard, G. S., Baker, D., Wolcott, M. J., Norwood, D. A. & Dauphin, L. A. Multi-platform comparison of ten commercial master mixes for probe-based real-time polymerase chain reaction detection of bioterrorism threat agents for surge preparedness. Forensic Sci. Int. 223, 292–297 (2012).

28. Ståhlberg, A., Kubista, M. & Pfaffl, M. Comparison of Reverse Transcriptases in Gene Expression Analysis. Clin. Chem. 50, 1678–1680 (2004).

29. Gutierrez, L., Mauriat, M., Pelloux, J., Bellini, C. & Van Wuytswinkel, O. Towards a Systematic Validation of References in Real-Time RT-PCR. Plant Cell 20, 1734–1735 (2008).

30. Remans, T. et al. Reliable Gene Expression Analysis by Reverse Transcription-Quantitative PCR: Reporting and Minimizing the Uncertainty in Data Accuracy. Plant Cell 26, 3829–3837 (2014).

31. Bustin, S. A. Why the need for qPCR publication guidelines?—The case for MIQE. Ongoing Evol. QPCR 50, 217–226 (2010).

32. Untergasser, A. et al. Disclosing quantitative RT-PCR raw data during manuscript submission: a call for action. Mol. Oncol. 17, 713–717 (2023).

33. Bustin, S. & Nolan, T. Talking the talk, but not walking the walk: RT-qPCR as a paradigm for the lack of reproducibility in molecular research. Eur. J. Clin. Invest. 47, 756–774 (2017).

34. Abdel Nour, A. M., Azhar, E., Damanhouri, G. & Bustin, S. A. Five Years MIQE Guidelines: The Case of the Arabian Countries. PLOS ONE 9, e88266 (2014).

35. Huggett, J., Dheda, K., Bustin, S. & Zumla, A. Real-time RT-PCR normalisation; strategies and considerations. Genes Immun. 6, 279–284 (2005).

36. Czechowski, T., Stitt, M., Altmann, T., Udvardi, M. K. & Scheible, W.-R. Genome-Wide Identification and Testing of Superior Reference Genes for Transcript Normalization in Arabidopsis. Plant Physiol. 139, 5–17 (2005).

37. Manoli, A., Sturaro, A., Trevisan, S., Quaggiotti, S. & Nonis, A. Evaluation of candidate reference genes for qPCR in maize. J. Plant Physiol. 169, 807–815 (2012).

38. Dekkers, B. J. W. et al. Identification of Reference Genes for RT–qPCR Expression Analysis in Arabidopsis and Tomato Seeds. Plant Cell Physiol. 53, 28–37 (2012).

39. Rácz, G. A., Nagy, N., Tóvári, J., Apáti, Á. & Vértessy, B. G. Identification of new reference genes with stable expression patterns for gene expression studies using human cancer and normal cell lines. Sci. Rep. 11, 19459 (2021).

40. Jo, J. et al. Conventionally used reference genes are not outstanding for normalization of gene expression in human cancer research. BMC Bioinformatics 20, 245 (2019).

41. Bas, A., Forsberg, G., Hammarström, S. & Hammarström, M.-L. Utility of the Housekeeping Genes 18S rRNA, β-Actin and Glyceraldehyde-3-Phosphate-Dehydrogenase for Normalization in Real-Time Quantitative Reverse Transcriptase-Polymerase Chain Reaction Analysis of Gene Expression in Human T Lymphocytes. Scand. J. Immunol. 59, 566–573 (2004).

42. Currie, K. GENORM: A generalized norm calculation. Comput. Geosci. 17, 77–89 (1991).

43. Andersen, C. L., Jensen, J. L. & Ørntoft, T. F. Normalization of Real-Time Quantitative Reverse Transcription-PCR Data: A Model-Based Variance Estimation Approach to Identify Genes Suited for Normalization, Applied to Bladder and Colon Cancer Data Sets. Cancer Res. 64, 5245–5250 (2004).

44. Pfaffl, M. W., Tichopad, A., Prgomet, C. & Neuvians, T. P. Determination of stable housekeeping genes, differentially regulated target genes and sample integrity: BestKeeper – Excel-based tool using pair-wise correlations. Biotechnol. Lett. 26, 509–515 (2004).

45. Xie, F., Xiao, P., Chen, D., Xu, L. & Zhang, B. miRDeepFinder: a miRNA analysis tool for deep sequencing of plant small RNAs. Plant Mol. Biol. 80, 75–84 (2012).

46. Lebuhn, M. et al. DNA and RNA extraction and quantitative real-time PCR-based assays for biogas biocenoses in an interlaboratory comparison. Bioengineering 3, 7 (2016).

47. Esteva-Socias, M., Gómez-Romano, F., Carrillo-Ávila, J. A., Sánchez-Navarro, A. L. & Villena, C. Impact of different stabilization methods on RT-qPCR results using human lung tissue samples. Sci. Rep. 10, 3579 (2020).

48. Bustin, S. A. & Nolan, T. Pitfalls of quantitative real-time reverse-transcription polymerase chain reaction. J. Biomol. Tech. JBT 15, 155 (2004).

49. Koppelkamm, A., Vennemann, B., Lutz-Bonengel, S., Fracasso, T. & Vennemann, M. RNA integrity in post-mortem samples: influencing parameters and implications on RT-qPCR assays. Int. J. Legal Med. 125, 573–580 (2011).

50. Vermeulen, J. et al. Measurable impact of RNA quality on gene expression results from quantitative PCR. Nucleic Acids Res. 39, e63–e63 (2011).

51. Die, J. V. & Román, B. RNA quality assessment: a view from plant qPCR studies. J. Exp. Bot. 63, 6069–6077 (2012).

52. Fleige, S. et al. Comparison of relative mRNA quantification models and the impact of RNA integrity in quantitative real-time RT-PCR. Biotechnol. Lett. 28, 1601–1613 (2006).

53. Angela Pérez-Novo, C., et al. Impact of RNA Quality on Reference Gene Expression Stability. BioTechniques 39, 52–56 (2005).

54. Bustin, S. Quantification of mRNA using real-time reverse transcription PCR (RT-PCR): trends and problems. J. Mol. Endocrinol. 29, 23–39 (2002).

55. Meijerink, J. et al. A Novel Method to Compensate for Different Amplification Efficiencies between Patient DNA Samples in Quantitative Real-Time PCR. J. Mol. Diagn. 3, 55–61 (2001).

56. Luu-The, V., Paquet, N., Calvo, E. & Cumps, J. Improved Real-Time RT-PCR Method for High-Throughput Measurements using Second Derivative Calculation and Double Correction. BioTechniques 38, 287–293 (2005).

57. Livak, K. J. & Schmittgen, T. D. Analysis of relative gene expression data using real-time quantitative PCR and the 2− ΔΔCT method. methods 25, 402–408 (2001).

58. Ruiz-Villalba, A., Ruijter, J. M. & van den Hoff, M. J. Use and misuse of Cq in qPCR data analysis and reporting. Life 11, 496 (2021).

59. Pfaffl, M. W. A new mathematical model for relative quantification in real-time RT–PCR. Nucleic Acids Res. 29, e45–e45 (2001).

60. Hellemans, J., Mortier, G., De Paepe, A., Speleman, F. & Vandesompele, J. qBase relative quantification framework and software for management and automated analysis of real-time quantitative PCR data. Genome Biol. 8, R19 (2007).

61. Pfaffl, M. W., Horgan, G. W. & Dempfle, L. Relative expression software tool (REST©) for group-wise comparison and statistical analysis of relative expression results in real-time PCR. Nucleic Acids Res. 30, e36–e36 (2002).

62. Lefever, S. et al. RDML: structured language and reporting guidelines for real-time quantitative PCR data. Nucleic Acids Res. 37, 2065–2069 (2009).

63. Ruijter, J. M. et al. RDML-Ninja and RDMLdb for standardized exchange of qPCR data. BMC Bioinformatics 16, 197 (2015).

64. Bustin, S. A. et al. The need for transparency and good practices in the qPCR literature. Nat. Methods 10, 1063–1067 (2013).

65. McGrath, J. C., Drummond, G., McLachlan, E., Kilkenny, C. & Wainwright, C. Guidelines for reporting experiments involving animals: the ARRIVE guidelines. Br. J. Pharmacol. 160, 1573–1576 (2010).

66. Bustin, S. A. et al. MIQE précis: Practical implementation of minimum standard guidelines for fluorescence-based quantitative real-time PCR experiments. BMC Mol. Biol. 11, 74 (2010).

67. Taylor, S. C. et al. The ultimate qPCR experiment: producing publication quality, reproducible data the first time. Trends Biotechnol. 37, 761–774 (2019).

68. Teytelman, L., Stoliartchouk, A., Kindler, L. & Hurwitz, B. L. Protocols.io: Virtual Communities for Protocol Development and Discussion. PLOS Biol. 14, e1002538 (2016).

69. LaFlamme M, Harney J, Hrynaszkiewicz I. A survey of researchers’ methods sharing practices and priorities. PeerJ (2024) doi:10.7717/peerj.16731.

70. S, B. L., et al. Promoting Reusable and Open Methods and Protocols (PRO-MaP). (2024) doi:10.2760/46124 (online),10.2760/58321 (print).

71. Schulz, R. et al. Is the future of peer review automated? BMC Res. Notes 15, 203 (2022).

72. Macleod, M., et al. The MDAR (Materials Design Analysis Reporting) Framework for transparent reporting in the life sciences. Proc. Natl. Acad. Sci. 118, e2103238118 (2021).

73. Weissgerber, T. et al. Automated screening of COVID-19 preprints: can we help authors to improve transparency and reproducibility? Nat. Med. 27, 6–7 (2021).

