## Supplement Table S1, S2 and Figure S1 for "Reporting quality of quantitative polymerase chain reaction (qPCR) methods in scientific publications"

**Table S1: Genetics and heredity article screening by journal**

| **Journal title** | **Articles screened**  n | **Articles included in the study**  n | **Articles included in the study**  n (%) |
| --- | --- | --- | --- |
| Nature Genetics | 30 | 6 | 20% |
| Genome Research | 36 | 1 | 2.77% |
| Molecular Biology and Evolution | 87 | 3 | 3.44% |
| Genome Biology | 51 | 7 | 13.72% |
| Genome Medicine | 33 | 1 | 3.03% |
| American Journal of Human Genetics | 35 | 6 | 17.14% |
| Genes Development | 7 | 3 | 42.85% |
| Molecular Therapy: The Journal of the American Society of Gene Therapy | 40 | 15 | 37.50% |
| Genetics in Medicine: Official Journal of the American College of Medical Genetics | 64 | 6 | 9.37% |
| Oncogene | 77 | 48 | 62.33% |
| American Journal of Medical Genetics C: Seminars in Medical Genetics | 12 | 0 | 0% |
| Genomics, Proteomics and Bioinformatics | 14 | 1 | 7.14% |
| Genomics | 47 | 14 | 29.78% |
| Molecular Autism | 11 | 0 | 0% |
| Human Genetics | 20 | 3 | 15% |
| NPJ Genomic Medicine | 17 | 4 | 23.52% |
| Horticulture Research* | 41 | 23 | 56.09% |
| The CRISPR Journal | 13 | 2 | 15.38% |
| PLoS Genetics | 89 | 28 | 31.46% |
| Human Molecular Genetics | 24 | 9 | 37.50% |

The following journals were excluded as they do not publish original research: Nature Reviews Genetics, Trends in Ecology and Evolution, Trends in Genetics, Annual Review of Genetics, Annual Review of Genomics and Human Genetics, Reviews in Mutation Research, and Current Opinion in Genetics and Development.

*Numbers presented in this table are higher than those in Table S2, as the genetics and heredity search included articles published in September and October 2021, whereas the search for plant sciences (Table S2) only included articles published in September 2021.

**Table S2: Plant sciences article screening by journal**

| **Journal title** | **Articles screened**  n | **Articles included in the study**  n | **Articles included in the study**  **n (%)** |
| --- | --- | --- | --- |
| Nature plants | 17 |  |  |
| Molecular plant | 20 | 7 | 35% |
| The plant cell | 17 | 8 | 47.05% |
| The new phytologist | 64 | 13 | 20.31% |
| Plant biotechnology journal | 18 | 9 | 50% |
| Plant physiology | 39 | 16 | 41.02% |
| Plant, cell and environment | 26 | 12 | 46.15% |
| The Plant journal: for cell and molecular biology | 39 | 19 | 48.71% |
| Journal of experimental botany | 52 | 22 | 40% |
| The Journal of ecology | 24 | 0 | 0% |
| Horticulture Research* | 20 | 9 | 45% |
| Journal of integrative plant biology | 11 | 3 | 27.27% |
| TAG. Theoretical and applied genetics | 25 | 6 | 24% |
| Frontiers in plant science | 288 | 84 | 29.16% |
| Molecular plant pathology | 11 | 9 | 81.81% |
| Phytomedicine: international journal of phytotherapy and phytopharmacology | 34 | 15 | 44.11% |
| Physiologia Plantarum | 17 | 0 | 0% |
| Plant & Cell Physiology | 15 | 5 | 33.33% |
| Environmental and Experimental Botany | 32 | 16 | 50% |

The following journals were excluded as they do not publish original research: Annual Review of Plant Biology, Trends in Plant Science, Annual Review of Phytopathology, Current Opinion in Plant Biology, Critical Reviews in Plant Sciences, and Phytochemistry Reviews. The journal Preslia was excluded as no articles were published in September 2021.

*All articles from Horticulture Research were also included in genetics and heredity (Table S1).

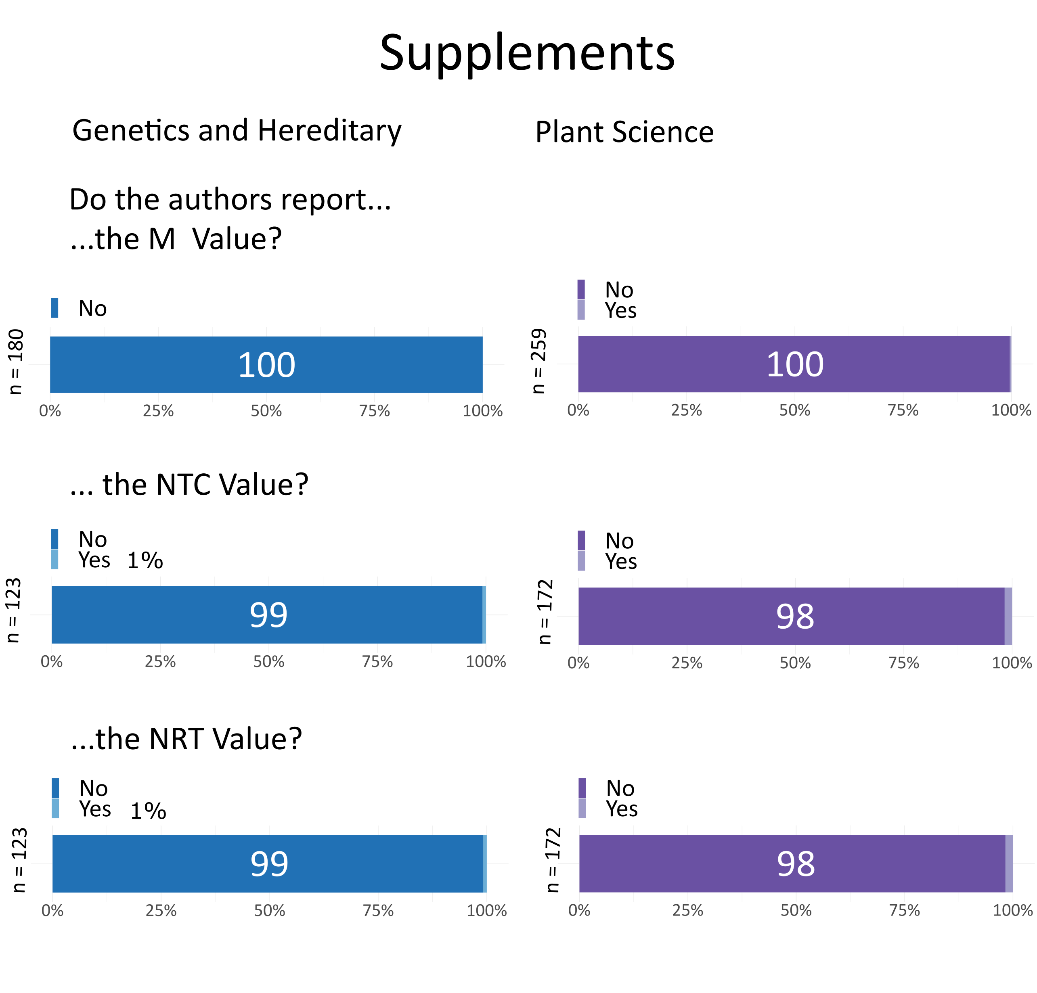

**Figure S1.** Lack of reporting of M values (control gene-stability measure), no template control (NTC) and no reverse transcriptase control (NRT).
